# Isolation and genomic analysis of secondary metabolism in cave Actinomycetota from biofilms and *Ceuthophilus*

**DOI:** 10.1101/2025.10.06.680698

**Authors:** Ilyana O. Bachmann, Brian O. Bachmann

## Abstract

We report the isolation and genomic sequencing of nine Actinomycetota obtained from both cave biofilms and trogloxenic *Ceuthophilus* (cave crickets). Strains were isolated from samples via actinomycetota-selective media and sequenced using nanopore sequencing. Provisional taxonomic inference of strains was performed by whole genome phylogenetic analysis, revealing a broad range of genera, with most strains clustering with known genera with digital DNA-DNA hybridization values less than 70%, and one strain not clustering with known genera. An analysis of secondary metabolism reveals the quantity and diversity of secondary metabolism in cave Actinomycetota.

## Description

The Actinomycetota phylum has been historically prolific in producers of bioactive secondary metabolites with clinical application (Yi, Shi, Cao, & Pan, 2025). Many antibiotics and anticancer drugs are bacterial secondary metabolites, derived from secondary metabolites, or address a target of a secondary metabolite (Newman & Cragg, 2020). For example, while first discovered in fungal strains, the β-lactam antibiotics, including penicillins, cephalosporins, clavams, and carbapenems, are also produced by various strains of Actinomycetota (Lewis et al., 2024). Anticancer secondary metabolites produced by Actinomycetota have roles in the treatment of a variety of cancer types. Daunomycin, used in the treatment of Acute Myeloid Leukemia and other cancer (Lewis et al., 2024), is produced by *Streptomyces* (Hutchinson, 1997). Carfilzomib, a derivative of epoxomicin, is produced by *Streptomyces* (Schorn et al., 2014). However, after decades of mining for bioactive secondary metabolites, the rates of discovery of novel compounds have slowed(Pye, Bertin, Lokey, Gerwick, & Linington, 2017).

Most Actinomycetota have been isolated from terrestrial ecosystems. The slowing rate of compound discovery is correlated with the slowing rate of discovery of novel Actinomycetota. It has been demonstrated that novel microbial taxa are more likely to produce novel chemistry (Yi et al., 2025). Moreover, metagenomic studies suggest a large reservoir of Actinomycetota remains to be cultivated (Bull & Goodfellow, 2019). Novel taxa can be discovered by developing new culture techniques or isolating strains from previously uninvestigated ecosystems.

In previous work, we have isolated novel bioactive secondary metabolites from cave actinomycetota (Covington, Spraggins, Ynigez-Gutierrez, Hylton, & Bachmann, 2018; Derewacz et al., 2014; Earl et al., 2018). Herein, we investigate the potential of cave ecosystems for the discovery of Actinomycetota strains with the potential to produce secondary metabolites. Our goal was to investigate two sources, cave microbial biofilms and trogloxenic insects, in particular cave crickets (*Ceuthophilus*), via Actinomycetota selective culture conditions, nanopore sequencing of isolated strains, and analysis of secondary metabolic biosynthetic gene clusters.

Two caves were targeted for this study. Hardin’s Cave, privately owned, and Snail Shell Cave, managed by the Southeastern Cave Conservancy. Hardin’s Cave is a 3.5-mile dry cave. Snail Shell cave comprises over nine miles of surveyed passages and is characterized by extensive flowing water passages, many of which require a boat for passage. Both cave ecosystems utilize allochthonous carbon input from the forest soil ecosystem. For this pilot assessment of hypogean taxonomic Collection of samples and crickets from caves. To collect samples from Snail Shell cave (Figure A), we developed a biofilm collection methodology that targeted biofilm from cave walls. Ceuthophilus samples were generated from whole crickets to encourage the isolation of spore-forming Actinomycetota. Three soil samples and three crickets were used in isolation workflows.

A wide variety of methods have been developed for the isolation of Actinomycetota from environmental samples. We selected two methods to enrich for the isolation of actinomycetes. As described in Methods, we adapted a standard soil dilution and plating methodology using humic acid vitamin (HV) agar with added antibiotics to suppress fungi and Gram-negative microorganism growth (Hayakawa & Nonomura, 1987). Second, we adapted a methodology to enrich the presence of motile spores (Hayakawa, Otoguro, Takeuchi, Yamazaki, & Iimura, 2000; Hop et al., 2011).

Colonies formed in HV agar primary plates, and fast-growing colonies were picked and subcultured in as little as one week (Figure B), and slow-growing colonies were subcultured in as much as three weeks. Subculture plates of International Streptomyces Protocol 1 (ISP1) medium agar, typtic soy broth (TSB) agar, and Bennett’s agar were streaked to subculture picked colonies from primary plates onto secondary plates. Photographs of isolated Actinomycetota are shown in Figure B. Single colonies were used to subculture onto tertiary plates of the best medium for growth, and these were used to inoculate liquid cultures in ISP1 or Bennett’s medium. From 25 mL cultures, we prepared glycerol stocks and used the majority for genomic sequencing.

Genomes of nine selected isolated strains sequenced by nanopore sequencing using Oxford Nanopore sequencing at Plasmidsaurus Inc. We evaluated two methodologies for sequencing samples. Genomic DNA from four liquid cultures was isolated using a classic protocol optimized for bulk isolation of genomic DNA from actinomycetes. Isolated DNA was of high purity (1.96-2.00 A_260_/A_280_, 1.97 -2.07 A_260_/A_230_) and submitted for sequencing. Alternatively, for six liquid cultures, we submitted cell pellets for automated DNA isolation using a ZymoBIOMICS™ 96 MagBead DNA Kit and nanopore sequencing. As shown in Table 1, we obtained high-quality assemblies for all samples. Notably, we obtained lower contig numbers and higher coverage from submitted cell pellets than from purified genomic DNA, though all genomes were of sufficient quality to permit taxonomic estimation and evaluation of secondary metabolic potential.

**Table 1.** List of strains isolated and sequenced in this study and taxonomic inference.

| Organism<sup>1</sup> | Total reads | Assembled coverage (fold) | Genome size (Mb) | Contigs | Genes annotated | Closest clustering species<sup>2</sup> | dDDH d<sub>4</sub> (%) | GenBank Accession |
| --- | --- | --- | --- | --- | --- | --- | --- | --- |
| IBHARD001 | 248,811 | 97 | 9.5 | 2 | 8,494 | *Nonomuraea muscovyensis* DSM 45913<br>*Planomonospora corallina* TBRC 4489 | 22.3<br>22.2 | PRJNA1308599 |
| IBHARD003 | 221,983 | 75 | 8.3 | 2 | 7,251 | *Streptomyces formicae* DSM 100524 | 28.6 | PRJNA1308605 |
| IBHARD004 | 380,186 | 99 | 6.8 | 1 | 6,546 | *Micromonospora kangleipakensis* DSM 45612 | 66.8 | PRJNA1308606 |
| IBHARD005 | 266,653 | 102 | 6.3 | 1 | 5,866 | *Nocardia soli* NBRC 100376 | 44.4 | PRJNA1308607 |
| IBSNAI012 | 352,887 | 102 | 6.6 | 1 | 6,126 | *Micromonospora palomenae* DSM 102131 | 68.5 | PRJNA1308591 |
| IBSNAI001 | 316,549 | 101 | 8.5 | 4 | 7,810 | *Streptomyces silvae* For3 | 69.7 | PRJNA1308129 |
| IBSNAI002 | 168,615 | 92 | 9.9 | 8 | 9,065 | *Streptomyces caledonius* MS1.HAVA.3 | 34.9 | PRJNA1308159 |
| IBSNAI004 | 69,803 | 45 | 9.4 | 9 | 8,707 | *Streptomyces kutzneri* DSM 40907 | 46.7 | PRJNA1308588 |
| IBSNAI006 | 378,350 | 98 | 9.9 | 2 | 8,943 | *Streptomyces lannensis* JCM16578 | 73.5 | PRJNA1308590 |

<sup>1</sup>IBHARD strains were isolated from Hardin's cave *Ceuthophilus* and IBSNAI strains from Snail Shell cave biofilms. <sup>2</sup>For the phylogenomic inference, all pairwise comparisons among the set of genomes were conducted using GBDP and accurate intergenomic distances inferred under the algorithm 'trimming' and distance formula $d_5$ . 100 distance replicates were calculated each. Digital DDH values and confidence intervals were calculated using the recommended settings of the GGDC 4.0.

Taxonomic estimation via whole gene taxonomic analysis using the Type (Strain) Genome Serve (TYGS), which estimates taxonomic assignments and phylogenies for submitted strains in comparison to a continuously updated database of type strains. Taxonomic distance is proportional in part to the Digital DNA-DNA Hybridization (sDDH) score shown, with all strains below the cutoff for preliminary designation as novel species. Figure C shows an example of phylogenetic analysis of strain IBHARD001, isolated from a cave cricket and analyzed by whole genome taxonomy using TYGS. This shows the tree inferred with FastME 2.1.6.1 (Lefort, Desper, & Gascuel, 2015) from GBDP distances calculated from genome sequences. The branch lengths are scaled in terms of GBDP distance formula *d*_*5*_ and indicate that this strain does not cluster with any of the species that it is most similar to, suggesting it may be a member of a previously undescribed genus. Future studies will be required to validate this preliminary assignment.

Secondary metabolic potential of genomically sequenced strains was performed using the antiSMASH program, which uses GenBank-formatted data to analyze for the presence of secondary metabolic biosynthetic gene clusters (BGCs). Gene clusters are classified into biosynthetic classes based on the predicted presence of diagnostic genes for each class. From antiSMASH analysis of the nine genomes, we tabulated the number of BGCs in four major classes. As shown in Panel D, isolates contain between 12 and 43 predicted BGCs, the number of which roughly scales with genome size. Number and types of biosynthetic classes varied between strains.

This pilot study defines the potential for the discovery of new taxa with secondary metabolic potential from cave ecosystems and trogloxenic insects. Future studies with a broader range of culture conditions and sample treatments can expand the scope of microorganismal discovery. Preliminary species and genera can be validated by follow-up studies. Promising strains containing biosynthetic gene clusters of interest can enter into a molecular discovery workflow to assess and validate the potential of novel cave taxa for generating novel bioactive secondary metabolites as leads for development in drug discovery programs.

## Data availability

GenBank accession numbers are provided in Table 1.

**Figure 1.**
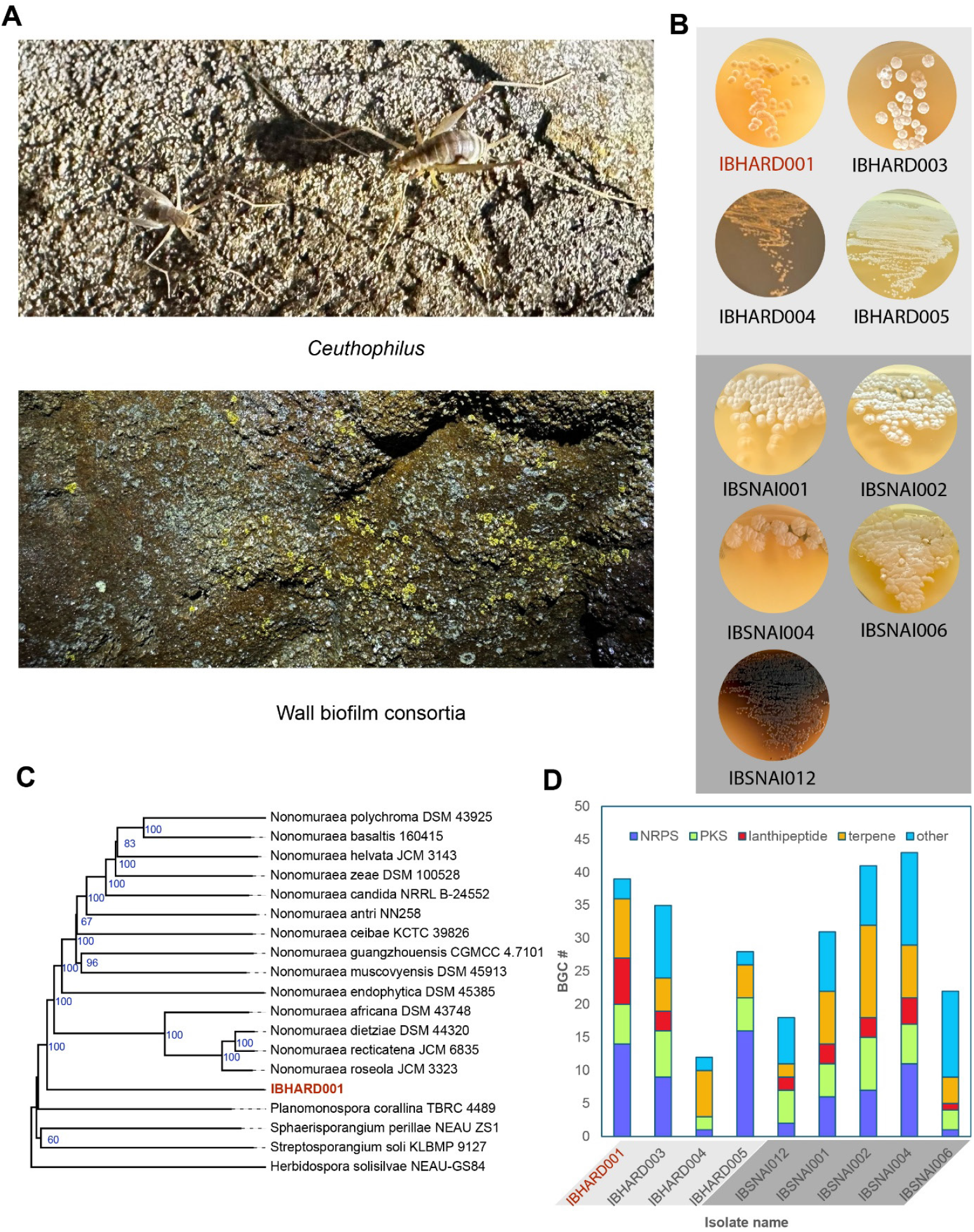
Isolation and characterization of hypogean Actinomycetota. **A**. *Ceuthophilus* and wall biofilms collected from expeditions for this study in Hardin’s Cave and Snail Shell Cave, respectively. **B**. Images of isolated strains growing on agar. **C**. Example of taxonomic estimation of IBHARD001, the most divergent actinomycete isolated in this study. Tree inferred with FastME 2.1.6.1 (Lefort et al., 2015) from GBDP distances calculated from genome sequences. The branch lengths are scaled in terms of the GBDP distance formula *d*_*5*_. The numbers on the above branches are GBDP pseudo-bootstrap support values > 60 % from 100 replications, with an average branch support of 85.8 %. The tree was rooted at the midpoint (Farris, 1972). **D**. Secondary metabolic potential quantified by antiSMASH (Blin et al., 2025). BGC for major classes are nonribosomal peptide synthetase (NRPS), polyketide synthase (PKS), lantipeptides, terpenes, and “others”.

## Methods

### Cave environmental sample collection

Sediment samples were collected from Snail Shell Cave, a hydrologically active cave located in Rockvale, Tennessee. The samples were collected immediately after the twilight zone, in the dark zone, from cave walls that demonstrated evidence of microbial colonization. A one cm^2^ sterile abrasive pad (Scotch-Brite) was used to collect biofilm sediments by scrubbing the wall in a circle with a ten cm diameter. Five sediment samples were collected and transported in sterile Falcon tubes, transferred into sterile petri dishes, and dried at 30 °C for two weeks. Dry weights were 215 - 553 mg. Sediment was removed from the pads aseptically by stretching and bending the pads. Cave-dwelling crickets were collected from Hardin’s Cave, located in Nashville, Tennessee. The crickets were captured in sealable plastic bags and crushed within ten minutes of capture. The crickets were then cut into fine pieces in a Petri dish, at 30 °C for one week to induce sporulation and further cut with a sterile razor blade to produce a coarse powder.

### Selective isolation method

Sediments and crickets were plated using two selective isolation methods. For both methods, the samples were first resuspended in 100 mL beakers (60 × 46 mm diameter) in 50 mL phosphate buffer (50 mM, pH = 7.0) containing 10 % soil extract, stirring with a glass rod for 30 seconds, followed by 90 minutes of rest at 30 °C. In the “motile spore preparation” method (Hayakawa et al., 2000; Hop et al., 2011), 8 mL of the upper portion of the sample solution was transferred into a 15 mL Falcon tube (16 × 105 mm). The tubes were centrifuged for 10 minutes at 1500 x g and allowed to rest at 30 ºC for 30 minutes to enable motile spores to circulate. From the middle of the volume, 3 mL was removed to a sterile Eppendorf tube and ten-fold serially diluted twice. In the “standard dilution method,” the above 90-minute rested solution was briefly stirred, allowed to settle for 30 seconds, and the suspension was diluted with sterile H_2_O, 1:100, 1:1K, and 1:10K dilutions.

All samples were plated on Humic Acid Vitamin agar (Hayakawa & Nonomura, 1987), ISP1, and Bennett’s Agar supplemented with a final concentration of nalidixic acid (20 mg/L), trimethoprim (10 mg/L), and cycloheximide (50 mg//L) to inhibit the growth of fungi and Gram-negative bacteria. The plates were incubated for 2 - 4 weeks at 30 ºC, and colonies of interest were picked and inoculated onto secondary purification plates by streaking onto ISP1 and Bennett’s media Agar. Individual colonies were used to streak tertiary isolation plates. Loops of mycelia from tertiary plates were used to inoculate ISP1 and Bennett’s liquid cultures (25 mL in a 250 mL Erlenmeyer flask). After 5-7 days of incubation with orbital shaking (230 rpm). A 0.5 mL portion of the culture was combined with an equal volume of sterilized 20% glycerol, 10% lactose, and stored at -80 °C. The remaining cultures were pelleted by centrifugation at 3348 x g for 20 minutes and stored at -80 ºC.

### Isolation of genomic DNA

Genomic DNA was isolated using a procedure adapted from standard protocols (Kieser, J., Buttner, Chater, & Hopwood, 2000). The frozen bacterial pellets were resuspended in 5 mL TE25S buffer. 100 µL of Lysozyme solution (100 mg mL^-1^ in nuclease-free H_2_O) and 200 µL/mL RNAse A (0.004 g in TE25S Buffer) were added. The sample was incubated for 60 minutes at 37 ºC, with occasional inversions. Then 50 µL Proteinase K solution (0.02 g in 1 mL H_2_O) was added and then mixed. 300 µL of SDS solution was added, and the sample was incubated for one hour at 55 ºC in a water bath, with occasional inversion. Then 1 mL of 5M NaCl and 650 µL CTAB/NaCl were added and mixed in, and the sample was incubated for 10 minutes at 55ºC. The samples were cooled to about 37 ºC, and 5 mL chloroform/isoamyl alcohol (24:1) was added and mixed via inversion for 30 minutes. The samples were centrifuged for 15 minutes at 3750 rpm, and the top layer was transferred to a fresh Falcon tube, being careful not to pull the interface layer. Then 0.6 volume of isopropanol was added and mixed by inversion. After about 3 minutes, the DNA was precipitated. Then the DNA was spooled with a glass hook and rinsed in 70% Ethanol. The DNA was transferred to a clean Eppendorf tube and air-dried for about 2.5 minutes. The DNA was then dissolved in 250 µL EB Buffer and heated to 55 ºC for 30 minutes. The DNA was incubated at four ºC for 3 days to fully rehydrate and homogenized by pipetting up and down using a 200 μL micropipette.

### Nanopore sequencing

Genomic DNA was isolated from IBSNAI001, IBSNAI002, IBSNAI004, and IBSNAI006 using a modified CTAB protocol from 25 mL cultures, which were diluted to 50 ng/μL. Pellets of IBHARD001, 003, 004, 005, and IBSNAI012 were generated from 25 mL liquid cultures. Bacterial Genome Sequencing for IBSNAI001, IBSNAI002, IBSNAI004, and IBSNAI006 was performed on isolated DNA by Plasmidsaurus Inc. using Oxford Nanopore Technology with custom analysis and annotation. Bacterial Genome Sequencing for IBHARD001, 002, 003, 004, 005, and for IBHARD012 was performed from pellets (30 - 50 mg), genomic DNA isolated using ZymoBIOMICS™ 96 MagBead DNA Kit (D4308/D4302) and sequenced as above.

### Taxonomic estimation

The genome sequence data were uploaded to the Type (Strain) Genome Server (TYGS), a free bioinformatics platform available under https://tygs.dsmz.de, for a whole genome-based taxonomic analysis (Meier-Kolthoff & Goker, 2019). The analysis also made use of recently introduced methodological updates and features. Information on nomenclature, synonymy, and associated taxonomic literature was provided by TYGS’s sister database, the List of Prokaryotic names with Standing in Nomenclature (LPSN, available at https://lpsn.dsmz.de) (Meier-Kolthoff, Carbasse, Peinado-Olarte, & Goker, 2022). The results were provided by the TYGS on 2025-08-11.

## Acknowledgements

Research was supported by NIH/NCI grant CA226833. We are grateful to Barry Walker, owner of Hardin’s Cave, and Southeastern Cave Conservancy, for access to Snail Shell Cave for academic research.

